# Disulfide-mediated tetramerization of TRAP1 fosters its antioxidant and pro-neoplastic activities

**DOI:** 10.1101/2024.09.19.613878

**Authors:** Fiorella Faienza, Claudio Laquatra, Matteo Castelli, Salvatore Rizza, Federica Guarra, Azam Roshani Dashtmian, Paola Giglio, Chiara Pecorari, Lavinia Ferrone, Elisabetta Moroni, Francesca Pacello, Andrea Battistoni, Giorgio Colombo, Andrea Rasola, Giuseppe Filomeni

**Affiliations:** Department of Biology, University of Rome Tor Vergata, Via della Ricerca Scientifica I-00133, Rome, Italy; Department of Biomedical Sciences, University of Padova, viale G. Colombo 3, I-35131, Padova, Italy; Department of Chemistry, University of Pavia, via Taramelli 12, I-27100, Pavia, Italy; Redox Biology, Danish Cancer Institute, Strandboulevarden 49, DK-2100, Copenhagen, Denmark; SCITEC-CNR, via Mario Bianco 9, I-20131, Milano, Italy

**Keywords:** oxidative stress, mitochondria, cysteine, tumorigenesis, metabolism

## Abstract

The mitochondrial chaperone TRAP1 exerts a protective function in cells exposed to diverse stress conditions in both physiological and pathological contexts. In cancer cells, it contributes to neoplastic progression ensuing metabolic rewiring and protection from oxidative insults.

TRAP1 works as a homodimer, but recent evidence has indicated that it can form tetramers whose functional effects remain elusive. Here, we find that TRAP1 forms redox-sensitive tetramers via disulfide bonds involving two critical cysteine residues, C261 and C573. TRAP1 tetramerization is elicited by oxidative stress and abrogated upon expression of the double C261S/C573R mutant. In cancer contexts, the expression of the TRAP1 C261S/C573R mutant is unable to inhibit the activity of its client succinate dehydrogenase and to confer protection against oxidative insults, and it hampers invasiveness of aggressive sarcoma cells.

Our data indicate that TRAP1 undergoes tetramerization in response to oxidative stress and identify C261 and C573 as critical for TRAP1 structural rearrangement and for functions.

## Introduction

The chaperone TRAP1, the mitochondrial paralog of HSP90, is a crucial homeostatic regulator. In cancer contexts, TRAP1 expression correlates with the acquisition of aggressive phenotypes, and its ablation or pharmacological inhibition hamper neoplastic growth (1). TRAP1 confers growth advantages in hypoxic environments and protection from the noxious effects of metabolic changes and from damage caused by reactive oxygen species (ROS) and redox dyshomeostasis (2—7). However, ROS have dichotomic effects in tumorigenesis as they might either induce cell death or activate signaling pathways sustaining malignancy (8). Hence, deciphering the molecular mechanisms and structural changes regulating TRAP1 activity is key to clarify the biological outputs of TRAP1 engagement during tumor progression and other pathophysiological conditions characterized by ROS dyshomeostasis.

TRAP1 activity dynamically adapts to fluctuations of intracellular conditions, and recent cryo-EM observations argue for TRAP1 undergoing quaternary structure changes as well. Particularly, it has been reported that TRAP1 may form tetramers as “dimers of dimers” with different conformations (9), an event which has been associated with perturbations of mitochondrial bioenergetics (10). However, it is unclear what triggers tetramer formation, what are the effects of tetramerization on TRAP1 chaperone activity, and if tetramers can regulate any specific biological function.

Here we demonstrate that TRAP1 can form tetramers following different oxidative stimuli via disulfide bridges involving C261 and C573, and that mutations of these residues affect TRAP1 activity and reduce its tumorigenic potential.

## Results and discussion

To address whether TRAP1 response to increased ROS levels is due to an intrinsic redox-sensitive nature of TRAP1 (i.e., its capacity to be direct target of ROS), we exposed a human recombinant TRAP1 to H_2_O_2_ and subjected it to non-reducing SDS-PAGE. In these conditions, we observed the appearance of a high molecular weight form of TRAP1, with an apparent molecular mass compatible with a tetramer, which disappeared upon incubation with the thiol-reducing agent DTT (**Fig. 1A)**. In order to determine which disulfide bridge(s) elicit TRAP1 tetramerization, we performed the same experiment after mutagenizing (C261, C501, C527, and C573). All four TRAP1 single mutants (i.e., C261S, C501S, C527A and C573R), as well as the C501S/C527A double mutant, were still able to form DTT-sensitive, high molecular weight structures upon H_2_O_2_ treatment. Vice-versa, the C261S/C573R TRAP1 mutant, as well as the cysteine-null (C_0_) variant, lost the ability to tetramerize (**Fig. 1B**), indicating that TRAP1 forms tetrameric structures via the engagement of disulfide bridges involving C261 and C573, as further confirmed by gel filtration chromatography (**Fig. 1C**).

**Figure 1.**
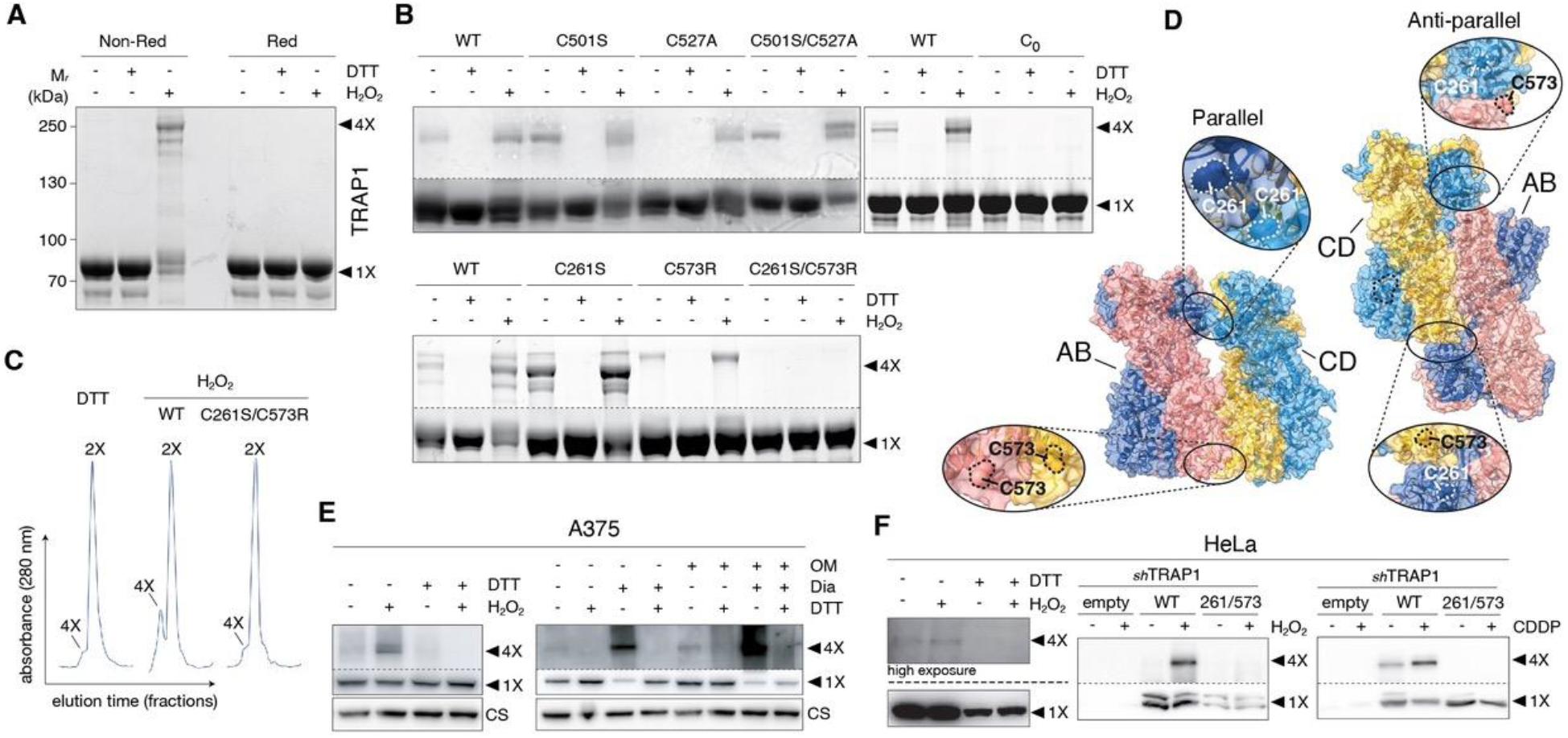
Recombinant TRAP1 (WT and cys-mutants) exposed to 100 μM H_2_O_2_ and/or 10 mM DTT for 30 min, and analyzed by (**A-B**) non-reducing or reducing SDS-PAGE and (**C**) gel-filtration chromatography. (**D**) Structural models of the parallel and antiparallel conformation of TRAP1 tetramers, as obtained from data-driven protein-protein docking simulations. Inset: magnification of C261 and C573-containing regions in TRAP1. Non-reducing Western blot analysis of TRAP1 in mitochondria isolated from (**E**) A375 cells treated with 100 μM H_2_O_2_ for 30 min; 1 mM diamide (Dia) for 30 min, or 10 μM oligomycin (OM) for 4 h, and (**F**) TRAP1-silenced (shTRAP1) HeLa cells expressing the WT or C261S/C573R HA-TRAP1 treated with H_2_O_2_ or 10 μM cisplatin (CDDP) for 24 h. DTT was used to reduce disulfides.

As these cysteines lie in distant regions of TRAP1, we speculated that potential disulfide bridges engaging these residues could link different TRAP1 dimers forming dimers-of-dimers. To investigate whether structural models of TRAP1 tetramers could be compatible with this hypothesis, we used a data-driven docking approach, defining the distance between C261 or C573 on one TRAP1 molecule and C261 or C573 on the other as restraints to guide protein-protein docking. We used structure 7KLV.pdb as the starting one for each dimer and obtained tetramer models compatible with the restraints in parallel or antiparallel configurations (**Fig. 1D**). The parallel configuration is compatible with the potential establishment of disulfide bonds between both C261, and both C573, of dimer AB and dimer CD. The antiparallel configuration indicates the possibility of forming disulfide bonds between C261 and C573 of different dimers (**Fig. 1D**).

To assess if TRAP1 oligomerization was also elicited in cells exposed to pro-oxidant conditions, we treated either human melanoma A375 cells or human cervix carcinoma HeLa cells with H_2_O_2_. Western blot analyses showed the presence of TRAP1 tetramers in both cell lines under these conditions (**Fig. 1E, F**). The monomer-to-tetramer shift was also induced by treatments with the thiol oxidant diamide and the ATP synthase inhibitor oligomycin, which also displayed a synergistic effect (**Fig. 1E**). In HeLa cells, the induction of TRAP1 tetramers upon H_2_O_2_ treatment was strongly enhanced by the ectopic expression of *wild-type* TRAP1 after stably knocking-down the endogenous protein, whereas the C261S/C573R mutant was unable to form tetramers (**Fig. 1F**). Similarly, the ROS-generating chemotherapeutic cisplatin (CDDP) prompted tetramer formation in cells harboring *wild-type* TRAP1, but not in those expressing the mutant (**Fig. 1F**). In each condition, the high molecular weight form of TRAP1 was not detectable after incubation with DTT (**Fig. 1E-F**), confirming that TRAP1 is a redox-sensitive protein that generates tetrameric structures via the engagement of disulfide bonds in response to a variety of oxidants.

We next investigated the functional effects of tetramers, i.e., whether they modify TRAP1 pro-neoplastic effects. To this aim, we studied a mouse malignant peripheral nerve sheath tumor cell model (sMPNST cells), as we had previously demonstrated TRAP1 involvement in shaping the tumorigenic features of this cancer type (11). We found that mouse sMPNST cells exhibited marked amounts of DTT-sensitive tetrameric TRAP1 even in basal conditions (**Fig. 2A, B**), and TRAP1 tetramerization was enhanced when cells were exposed to H_2_O_2_ (**Fig. 2A**). Similarly, we observed DTT-sensitive tetramers after re-expressing *wild-type* human TRAP1 in cells where the endogenous one had been knocked-out, whereas we could not detect any tetramers in cells reconstituted with the C261S/C573R TRAP1 mutant, even upon treatment with H_2_O_2_ or diamide (**Fig. 2B, C**). Expression of this mutant markedly hampered the *in vitro* tumorigenicity of sMPNST cells, as they displayed a limited capacity to form foci and to grow in 3D, both as spheroids and in branching morphogenesis experiments (**Fig. 2D, E**), but had no influence on the proliferation rate (not shown). Taken together, these observations indicate that the expression of the C261S/C573R mutant hinders anoikis-resistant growth and affects invasiveness of cancer cells.

**Figure 2.**
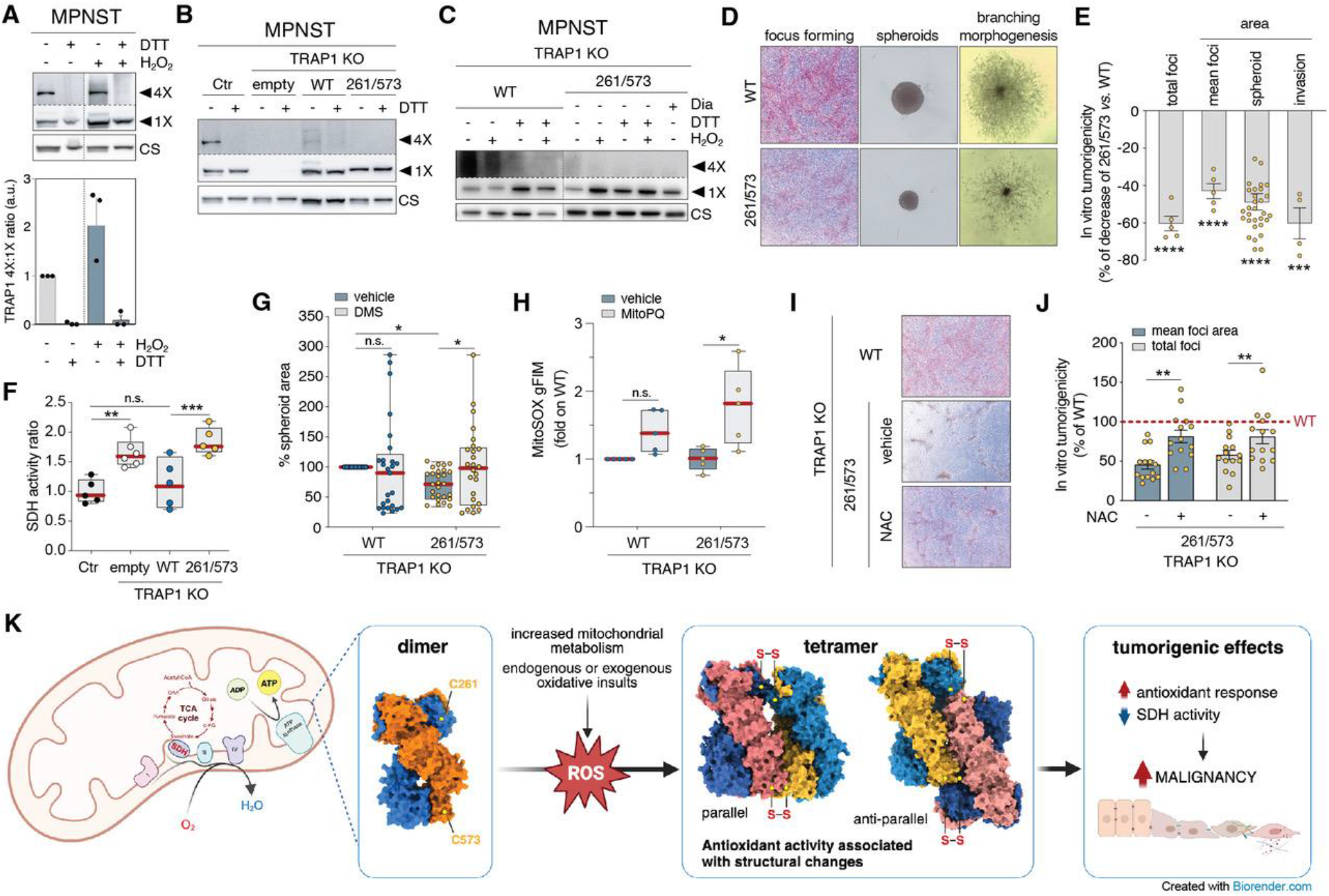
Non-reducing Western blot analysis of TRAP1 in: (**A**) mitochondria isolated from sMPNST cells treated with H_2_O_2_; (**B**) TRAP1-KO sMPNST cells transfected with HA-TRAP1 *wild-type*, C261S/C573R mutant, or an empty vector; (**C**) Same cells as in B treated with 1 mM diamide (Dia) or 100 μM H_2_O_2_ for 30 min. DTT was used to reduce disulfides. Densitometry of the ratio tetrameric (4X) *vs*. monomeric (1X) TRAP1 band intensities normalized on citrate synthase is shown at the bottom of A. (**D**) Representative images of tumor foci, spheroids and invasive masses of TRAP1-KO sMPNST cells reconstituted as in B. (**E**) Evaluation of total foci (n=5; ****, *p*<0.0001); foci area (n=5; ****, *p*<0.0001); spheroid size (n=30; ****, *p*<0.0001), and invasive area (n=4; ***, *p*<0.001) by ImageJ software. Data are reported as % of decrease of these phenotypes in TRAP1-C261S/C573R versus TRAP1-WT-expressing cells and expressed and mean ± S.D. Same cells as in B were used to assess: (**F**) succinate dehydrogenase (SDH) activity (**, *p*<0.01; ****, *p*<0.0001; *n*.*s*., not significant) and (**G**) spheroid area upon treatment with 2 mM dimethyl succinate (DMS) for 10 days (n=25; *, *p*<0.05; *n*.*s*., not significant — upon *t*-test analysis). (**H**) ROS levels evaluated upon treatment with 1 μM mito-paraquat (mito-PQ) for 2 h by MitoSOX staining. Data are reported as fluorescence intensity mean (n=5; *, *p*<0.05). **I**) Representative images of tumor foci of TRAP1-KO sMPNST cells reconstituted as in B, treated with 5 mM *N*-acetyl-L-cysteine (NAC) every 2 days for 10 days. **J**) Evaluation of total foci (n=14; **, *p*<0.01), foci area (n=14; **, *p*<0.01), by ImageJ software. Data are reported as % respect to WT and expressed and mean ± S.D. **K**) Schematic model for TRAP1 tetramers formation and tumorigenetic effects. Two-way ANOVA was applied for all statistical analyses, unless otherwise indicated..

We previously demonstrated that TRAP1 enhances tumor progression by downregulating SDH activity in neoplastic cells, thus inducing an accumulation of the oncometabolite succinate (3). We then evaluated SDH activity as a *bona fide* measurement of TRAP1 chaperone function. We found that expression of the C261S/C573R TRAP1 mutant almost doubled SDH activity with respect to sMPNST cells carrying *wild-type* TRAP1, thus functionally mimicking its ablation (**Fig. 2F**). Moreover, cell treatment with a membrane-permeable succinate analogue rescued the ability of C261S/C573R mutant-expressing sMPNST cells to form spheroids of the same size of *wild-type* cells (**Fig. 2G**), highlighting the importance of TRAP1-dependent modulation of SDH activity for tumorigenicity.

As TRAP1 can shield cells from oxidative stress (12), we investigated whether tetramer formation could play a role in its antioxidant function. We found that sMPNST cells treated with the redox cycler mito-paraquat (mitoPQ) underwent, as expected, a marked increase in mitochondrial superoxide levels. However, oxidative stress was much higher when TRAP1-KO cells were reconstituted with the C261S/C573R mutant than with *wild-type* TRAP1 (**Fig. 2H**). In agreement with the evidence that several neoplastic models require an increased antioxidant capacity for sustaining tumor progression (4), we observed that expression of the C261S/C573R mutant hampered the focus forming ability of sMPNST cells, which was almost fully restored by incubation with the thiol reducing agent *N*-acetyl-L-cysteine (NAC) (**Fig. 2I, J**). These results provide clear evidence that TRAP1 forms tetramers under oxidative conditions *via* the formation of disulfide bonds. Although qualitative, our structural models are compatible with (and reminiscent of) the experimental structures from Agard’s group (9).

The identification of the C261 and C573 as the residues responsible for TRAP1 tetramerization provides the first example of a detailed structure-function relationship that connects chaperone activity to cancer cell biology and suggests that the use of new classes of Cys-targeting drugs (as proposed in 13) might disclose novel and highly selective therapeutic opportunities by specifically affecting subsets of TRAP1 activities and interactions.

Overall, our findings argue for the formation of higher order structures being a general mechanism by which TRAP1 shields cells from noxious stimuli and suggest that such structures could expand the number of TRAP1 clients, facilitating its interactions with high molecular weight protein complexes. This model resembles the one already described for peroxiredoxins (Prxs) (14) and proposes TRAP1 as a novel mitochondrial “redox sensor”.

## Materials and Methods

Recombinant TRAP1 was generated as previously described (15) and treated with different oxidant species at room temperature. TRAP1-deficient human and murine tumor cells were reconstituted with WT and C261/C573R mutants upon transfection with Lipofectamine 3000. Tumorigenicity, SDH activity and ROS were assessed as previously reported (3). Models of TRAP1 tetramers were obtained out using PIPER in Schrodinger Maestro suite 2022-4 (www.schrodinger.com) with different constrains applied. See SI Appendix for detailed methods.

## Acknowledgments

This work was supported by: the Danish Cancer Society (KBVU R146-A9414, R231-A13855, and R352-A20537 to G.F.); the Novo Nordisk Foundation (NNF18OC0052550 and NNF22OC0079352 to G.F.); the Italian Association for Cancer Research (IG2022-27139 to G.C.; IG2023-29221 to G.F; IG2023-29351 to A.R.; Individual Fellowship Love Design 2021—ID 26647-2021 to F.G.; AIRC fellowship for Italy to F.F.); the Italian Ministry of University and Research (Progetti di Ricerca di Rilevante Interesse Nazionale PRIN MUR 2022C423E7 to A.R. and G.F., and MUR 20209KYCH9 to A.R.); Padova University (BIRD project 231497 to A.R.).

## Supplementary information

### Methods

#### Plasmids

TRAP1 WT and redox mutants (C501S, C527A, C501S/C527A, C261S, C573R, C261S/C573R) were cloned in pET-26b(+) plasmid for bacterial expression and in pcDNA3.1(+)-C-HA plasmid for mammalian expression using the Gene Synthesis & DNA Synthesis service from GenScript Biotech. TRAP1 WT in pBABE vector was mutagenized to produce redox mutants (C261S, C573R, C261S/C573R) using the following primers: C261S 5’-CACCTGAAATCCGACTCCAAGGAGTTTTCCAGC-3’, C573R 5’-CAGGTCCCCAGCCGCCGAGCGCCTATCAGAGAA-3’. All pBABE plasmids were used for viruses production.

#### Production of recombinant proteins

Recombinant TRAP1 (WT and redox mutants) was produced in BL21(DE3) *E. coli* cells after 3 h-induction with 1 mM IPTG (Isopropil-β-D-tiogalattopiranoside, VWR). Protein purifications were performed using Ni-NTA resin (Quiagen) according to manufacturer’s instructions. Proteins release from the resin packed in a FPLC column was achieved using a linear imidazole gradient generated by mixing *buffer A* (50 mM Phosphate Buffer, 250 mM NaCl, 10 mM imidazole, 10 mM β-mercaptoethanol, pH 7.8) and *buffer B* (buffer A containing 250 mM imidazole). Eluted fractions containing recombinant proteins were then collected and combined. After purification, the proteins were dialyzed in the *storage buffer* (50 mM Tris-HCl, 150 mM NaCl, 1 mm DTT, 1 mM EDTA, pH 7.5) and stored into aliquots.

#### Redox treatments

Purified TRAP1 (WT and redox mutants) were dialyzed in *storage buffer*. Before treatments, DTT was removed by Zeba Spin Desalting Columns (Thermo Fisher Scientific) and proteins transferred in a *reaction buffer* (150 mM NaCl, 50 mM TRIS, pH 7.5). Oxidation and reduction were performed at 37 °C by incubations with 100 μM H_2_O_2_ (Sigma Aldrich) and 50 mM DTT (Sigma Aldrich), respectively, for 30 min.

#### Non-reducing SDS-PAGE

After treatments with oxidants or reductants, proteins were denatured with sample buffer (NuPAGE LSD sample buffer, Thermo Fisher Scientific), with or without the reducing agent (NuPAGE Sample reducing agent, Thermo Fisher Scientific), and loaded on gel for SDS-PAGE. Proteins were visualized by Coomassie Brilliant Blue staining (Sigma Aldrich) or with Stain-free technology (Bio-Rad Laboratories).

#### Gel filtration chromatography

The quaternary structure of WT and C261S/C573R mutant TRAP1 protein was analyzed by size-exclusion chromatography using the prepacked column Superdex 200 HR 10/30 (Amersham Pharmacia Biotech). Before loading, the buffer was changed by Zeba Spin Desalting Columns (Thermo Fisher Scientific) to remove DTT. One hundred and fifty mg of each protein sample was applied to the column and eluted with 20 mM TRIS, 150 mM NaCl, pH 7. Fractions of 0.5 ml were collected and analyzed by SDS-PAGE to verify that all the observed elution peaks were due to TRAP1. The apparent molecular weight of the different forms of TRAP1 was determined using a calibration line generated by eluting standard proteins of known molecular weight in the same buffer.

To study the effect of the oxidant treatment on the quaternary structure of WT and cysteine mutants of TRAP1, each protein was incubated with 100 μM hydrogen peroxide and subjected to gel filtration.

#### Structural models

Prior to docking calculations, the Cryo-EM structure of tetrameric TRAP1 (PDB: 7KLV(https://doi.org/10.2210/pdb7KLV/pdb)) was subjected to preparation steps. The structure was preprocessed with protein preparation wizard tool in Maestro. Missing loops were reconstructed accordingly to UniProt Q12931 with crosslink in Schrodinger Maestro suite 2022-4 (www.schrodinger.com). Missing loops in protomer A were residues 335-360, 558-573 and 631-637. Missing loops in protomer B were residues 357-361, 558-573 and 625-630. Subsequently, the resulting loops underwent a minimization step with Prime until the RMS gradient convergence of 0.01 kcal/mol/Å was reached (solvation model: VSGB; Force field: OPLS4).

Protein-protein docking was carried out using PIPER in Schrodinger Maestro suite 2022-4 (www.schrodinger.com). Distance constraints were used as follows:

- Parallel configuration was obtained with C261(protomer-A)-C261(protomer-D) and C573(protomer-B)-C573(protomer-C) constraints.
- Antiparallel configuration was obtained with C261(protomer-A)-C573(protomer-C) and C261(protomer-D)-C573(protomer-B) constraints.

For each configuration, the best pose was optimized through minimization in explicit solvent (TIP3P water) with Prime (100ps). The last frame of minimization was used as starting conformation for 10 ns of MD (without any constraints) using the Desmond engine with default parameters (**1**), and TIP3P water as a solvent model (**2**). The last frame of the MD simulations is shown in **Fig. 1D**.

#### Cell lines

A375 and HeLa cell lines were obtained from the American Type Culture Collection (ATCC, Virginia, USA). Malignant peripheral nerve sheath tumor cells (sMPNST) were established from neurofibromin 1 (Nf1)-deficient skin precursors (SKP) and were kindly provided by Dr. Lu Q. Le, University of Texas Southwestern Medical Center, Dallas, TX. Cells were grown in Dulbecco’s Modified Eagle’s Medium (DMEM) and RPMI-1640 supplemented with 10% fetal bovine serum (FBS, Thermo Fisher Scientific) 100 U/ml and 1% penicillin/streptomycin (Euroclone). All cells were cultured at 37°C in a humidified atmosphere containing 5% CO_2_.

#### Generation of sMPNST mutant cells

pBABE vectors containing both the WT and the C261S/C573R form of TRAP1 were used to stably transfect sMPNST TRAP1 KO cells previously generated by Sanchez-Martin et al. (**3**). pBABE vectors were co-transfected with packaging plasmids PMD2.G and psPAX2 into HEK 293 cells for viral production. Recombinant virus was used to infect sMPNST cells that were subsequently selected with 0.75 mg/ml G418 (Sigma Aldrich).

#### Generation of TRAP1-depleted HeLa cells

Stable TRAP1-depleted HeLa cells were generated by transfection of a TRAP1 3’UTR-targeting pre-miRNA-containing plasmid as previously described (**4**). The plasmid was generated by cloning the following constructs in pcDNA6.2-GW/EmGFP-miR vector using the BLOCK-iT™ Pol II miR RNAi Expression Vector Kit (Life Technologies): Top Strand: 5’-TGCTGAGGTAAATAAAGCTCAAGGAGGTTTTGGCCACTGACTGACCTCCTTGATTTATTTACCT-3’; Bottom Strand: 5’-CCTGAGGTAAATAAATCAAGGAGGTCAGTCAGTGGCCAAAACCTCCTTGAGCTTTATTTACCTC-3’. Upon transfection, cells were cultured for 4 weeks in the presence of Blasticidin S and GFP-expressing cells were enriched twice using BD FACSMelody Cell Sorter (Beckman Coulter).

#### Cell treatments

For redox treatments cells were incubated with 100 μM H_2_O_2_ or with 50 mM DTT for 10 min in PBS at room temperature. Other treatments were performed as follow: a) diamide (Sigma Aldrich) was used 1 mM for 30 min; b) oligomycin A (Calbiochem) was used 10 μM for 4 h; c) cisplatin (CDDP, Sigma Aldrich) was used 10 μM for 24 h; d) dimethylsuccinate (DMS, SAFC) was used 2 mM for 10 days; e) MitoPQ (Cayman Chemical) was used 1 μM for 2 h; *N*-acetyl-L-cysteine (NAC, Sigma Aldrich) was used 5 mM every 2 days for 10 days.

#### Mitochondrial isolation

Mitochondria were isolated upon cell disruption with a glass-Teflon potter in RLM buffer composed of 250 mM sucrose, 10 mM Tris-HCl, 0.1 mM EGTA-Tris, pH 7.4. Sixty μg of mitochondria were resuspended in RLM buffer and treated as described above (*Cell treatments*). After treatments, mitochondria were pulled down at 8,200 rpm for 30 min, lysed in EB buffer (composed of NaCl 150 mM, Tris 20 mM pH 7.4, EDTA 5 mM, 1% Triton X-100, Glycerol 10%) and incubated on ice for 20 min and clarified at 18,000 x *g*. Supernatant was collected and used for Western blot analyses.

#### Western Blot analyses

Cells were lysed with Cell Lytic (Sigma Aldrich) plus protease inhibitors (Protease Inhibitors Cocktail, Sigma-Aldrich) and phosphatases inhibitors (1 mM Sodium fluoride and 1 mM Sodium orthovanadate, Sigma-Aldrich). After clarification protein extracts were quantified with DC protein assay (Bio-Rad Laboratories) protocol and then processed as described above (*Non reducing SDS-PAGE*). After SDS-PAGE, protein complexes were transferred onto nitrocellulose or PVDF membranes and probed with anti-TRAP1 (anti-murine TRAP1 BD and anti-human TRAP1 SC-73604 and SC-13557) or anti-citrate synthase (Abcam ab96600) antibodies.

#### Measurement of succinate:coenzyme Q reductase (SQR) activity of SDH

Succinate dehydrogenase (SDH) activity was measured as already described by Sanchez-Martin et al. (**3**) Briefly, samples were collected and lysed at 4°C in a buffer composed of 25 mM potassium phosphate, pH 7.2, and 5 mM MgCl_2_ containing protease and phosphatase inhibitors. Total lysate was quantified with BCA protein assay Kit (Thermo-Scientific), and 40 μg of protein per trace was incubated 10 min at 30 °C in the presence of 20 mM sodium succinate and 10 mM alamethicin. After incubation, a mix composed of 5 mM sodium azide, 5 μM antimycin A, 2 μM rotenone, and 65 μM Coenzyme Q1 was added. SDH activity was measured by following the reduction of 2,6-dichlorophenolindophenol (DCPIP) at 600 nm (ε = 19.1 nM^−1^ cm^−1^) at 30°C. Each measurement was normalized for protein amount.

#### Measurement of ROS levels

ROS levels were evaluated on sMPNST KO cells expressing the WT and C261S-C573R variants of TRAP1 treated or not with mitoPQ 1μM for 2 h. Upon treatment, cells were stained for 30 min with 5 μM MitoSOX and analyzed with a LSRFortessa X-20 flow cytometer (Becton Dickinson).

#### *In vitro* tumorigenic assay

Focus forming assays were performed on sMPNST cells grown in 12-well culture plates in DMEM medium supplemented with 10% fetal bovine serum. When cells reached sub-confluence, serum concentration was decreased to 1%. After 10 days, plates were washed in PBS, fixed in methanol for 30 min, stained with GIEMSA solution for 1 h and analyzed with ImageJ software.

#### Spheroids formation and in vitro migration assay

For spheroid generation, 5000 sMPNST cells were plated in round-bottom 96 well plates, centrifuged at 100 x *g* for 1 min at room temperature and incubated at 37°C 5% CO_2_. After 48 h, 50 μl of DMEM was replaced with same volume of Matrigel, centrifuged at 100 x *g* for 1 min at 4°C, and incubated at 37°C, 5% CO2. After 5 days, cell spreading was analyzed with ImageJ software.

## References

1. I. Masgras et al., Tumor growth of neurofibromin-deficient cells is driven by decreased respiration and hampered by NAD+ and SIRT3. Cell Death Differ. 29, 1996–2008 (2022).

2. S. Yoshida et al., Molecular chaperone TRAP1 regulates a metabolic switch between mitochondrial respiration and aerobic glycolysis. Proc. Natl. Acad. Sci. USA. 110, E1604–12 (2013).

3. M. Sciacovelli et al., The mitochondrial chaperone TRAP1 promotes neoplastic growth by inhibiting succinate dehydrogenase. Cell Metab. 17, 988–999 (2016).

4. G. Guzzo, M. Sciacovelli, P. Bernardi P, A. Rasola, Inhibition of succinate dehydrogenase by the mitochondrial chaperone TRAP1 has anti-oxidant and anti-apoptotic effects on tumor cells. Oncotarget. 5, 11897–908 (2014).

5. G. Cannino et al., The mitochondrial chaperone TRAP1 regulates F-ATP synthase channel formation. Cell Death Differ. 29, 2335–2346 (2022).

6. I. Masgras et al., The molecular chaperone TRAP1 in cancer: From the basics of biology to pharmacological targeting. Semin. Cancer Biol. 76, 45–53 (2021).

7. A. Rasola, L. Neckers, D. Picard, Mitochondrial oxidative phosphorylation TRAP(1)ped in tumor cells. Trends Cell Biol. 24, 455–63 (2014).

8. E.C. Cheung, K.H. Vousden, The role of ROS in tumour development and progression. Nat. Rev. Cancer. 22, 280–297 (2022).

9. Y. Liu et al., Cryo-EM analysis of human mitochondrial Hsp90 in multiple tetrameric states. bioRxiv 2020.11.04.368837 (2020).

10. A. Joshi et al., The mitochondrial HSP90 paralog TRAP1 forms an OXPHOS-regulated tetramer and is involved in mitochondrial metabolic homeostasis. BMC Biol. 18, 10 (2020)

11. I. Masgras et al., Absence of Neurofibromin Induces an Oncogenic Metabolic Switch via Mitochondrial ERK-Mediated Phosphorylation of the Chaperone TRAP1. Cell Rep. 18, 659–672 (2017).

12. G. Hua, Q. Zhang, Z. Fan, Heat shock protein 75 (TRAP1) antagonizes reactive oxygen species generation and protects cells from granzyme M-mediated apoptosis. J. Biol. Chem 282, 20553–20560 (2007)

13. M. Takahashi et al., DrugMap: A quantitative pan-cancer analysis of cysteine ligandability. Cell. 187, 2536-2556.e30 (2024).

14. Z.A. Wood, E. Schröde J. Robin Harris, L.B. Poole, Structure, mechanism and regulation of peroxiredoxins. Trends Biochem. Sci. 28, 32–40 (2003).

15. E. Papaleo et al., TRAP1 S-nitrosylation as a model of population-shift mechanism to study the effects of nitric oxide on redox-sensitive oncoproteins. Cell Death Dis. 14, 284 (2023)

## References to Methods

1. K.J. Bowers et al., Scalable algorithms for molecular dynamics simulations on commodity clusters. In: Proceedings of the 2006 ACM/IEEE conference on Supercomputing (SC ‘06). Association for Computing Machinery, New York, NY, USA, 84–es.

2. W.L., Jorgensen et al., Comparison of simple potential functions for simulating liquid water. J. Chem. Phys. 79, 926–935 (1983).

3. C. Sanchez-Martin et al., Rational Design of Allosteric and Selective Inhibitors of the Molecular Chaperone TRAP1. Cell Rep. 31, 107531 (2020).

4. S. Rizza et al., S-nitrosylation of the Mitochondrial Chaperone TRAP1 Sensitizes Hepatocellular Carcinoma Cells to Inhibitors of Succinate Dehydrogenase. Cancer Res. 76, 4170–4182 (2016).

